# Phosphoproteomic Identification of Vasopressin-Regulated Protein Kinases in Collecting Duct Cells

**DOI:** 10.1101/2020.07.22.215897

**Authors:** Arnab Datta, Chin-Rang Yang, Karim Salhadar, Chung-Lin Chou, Viswanathan Raghuram, Mark A. Knepper

**Affiliations:** Epithelial Systems Biology Laboratory, Systems Biology Center, National Heart, Lung, and Blood Institute, National Institutes of Health, Bethesda, Maryland; Yenepoya Research Center, Yenepoya (Deemed to be University), University Road, Deralakatte, Mangalore 575018, Karnataka, India

**Keywords:** mpkCCD, GPCR signaling, V2 receptor signaling, Desmopressin

## Abstract

**Background and Purpose:** The peptide hormone vasopressin regulates water transport in the renal collecting duct largely via the V2 receptor, which triggers a cAMP-mediated activation of a protein kinase A (PKA)-dependent signaling network. The protein kinases downstream from PKA have not been fully identified or mapped to regulated phosphoproteins.

**Experimental Approach:** We carried out systems-level analysis of large-scale phosphoproteomic data quantifying vasopressin-induced changes in phosphorylation in aquaporin-2-expressing cultured collecting duct cells (mpkCCD). Quantification was done using stable isotope labeling (SILAC method).

**Key Results:** 9640 phosphopeptides were quantified. Stringent statistical analysis identified significant changes in response to vasopressin in 429 of these phosphopeptides. The corresponding phosphoproteins were mapped to known vasopressin-regulated cellular processes. The vasopressin-regulated sites were classified according to the sequences surrounding the phosphorylated amino acids giving 11 groups distinguished predominantly by the amino acids at positions +1, −3, −2 and −5 relative to the phosphorylated amino acid. Among the vasopressin-regulated phosphoproteins were 25 distinct protein kinases. Among these, six of them plus PKA appeared to account for phosphorylation of more than 80% of the 313 vasopressin-regulated phosphorylation sites. The six downstream kinases were salt-inducible kinase 2 (Sik2), cyclin-dependent kinase 18 (PCTAIRE-3, Cdk18), calmodulin-dependent kinase kinase 2 (Camkk2), protein kinase D2 (Prkd2), mitogen-activated kinase 3 (ERK1; Mapk3), and myosin light chain kinase (Mylk).

**Conclusion and Implications:** In V2 receptor-mediated signaling, PKA is at the head of a complex network that includes at least 6 downstream vasopressin-regulated protein kinases that are prime targets for future study. The extensive phosphoproteomic data generated in this study is provided as a web-based data resource for future studies of G-protein coupled receptors.

## INTRODUCTION

G-protein coupled receptors (GPCRs) are central to the regulation of a variety of physiological processes and are frequently targeted for pharmacological treatments (1). One such GPCR is the vasopressin V2 receptor, expressed in renal collecting duct cells where it regulates the transport of water, sodium and urea (2). Signaling through the V2 receptor controls water transport through effects on the water channel, aquaporin-2 (AQP2). With vasopressin binding, the heterotrimeric G-protein α subunit, G_α_s, activates adenylyl cyclase 6 (3) and increases cAMP production. Loss of this signaling pathway results in diabetes insipidus. Central diabetes insipidus is treated with a V2 receptor-selective vasopressin analog dDAVP (1-deamino-8-D-arginine vasopressin), often referred to as ‘desmopressin’ (4). Defects in this signaling pathway can also result in abnormal renal water retention, e.g. the syndrome of inappropriate antidiuresis (SIADH), characterized by dilutional hyponatremia. Hyponatremia of all causes accounts for as much as 30% of hospitalized patients in tertiary care centers (5). Treatment employs a drug class known as ‘vaptans’ (e.g. tolvaptan), which function as V2 receptor antagonists (6). The clinical use of vaptans has recently expanded because of FDA-approval of vaptans in autosomal dominant polycystic kidney disease (7). In previous studies, we have modeled SIADH by water loading in rodents infused with dDAVP (8).

The functional effects of vasopressin at a cellular level in collecting duct principal cells are shown in Figure 1, which summarizes an extensive literature on the topic (9–33). Most of these downstream effects are believed to be mediated by protein kinase A (PKA) (34) (Figure 1). The details of vasopressin signaling remain incomplete. Here, we carried out systems-level analysis of large-scale phosphoproteomic data quantifying vasopressin-induced changes in phosphorylation in AQP2-expressing cultured mpkCCD cells. The data were used (a) to construct a publicly accessible web resource useful for the study of signaling by Gαs-coupled GPCRs; and (b) to identify a set of protein kinases downstream from PKA that are likely responsible for many of the phosphorylation changes triggered by vasopressin in collecting duct cells. We propose that the new information presented in this paper will be useful, not only for understanding water balance disorders, but also for understanding regulation by other G_α_s-coupled receptors in other tissues.

**Figure 1.**
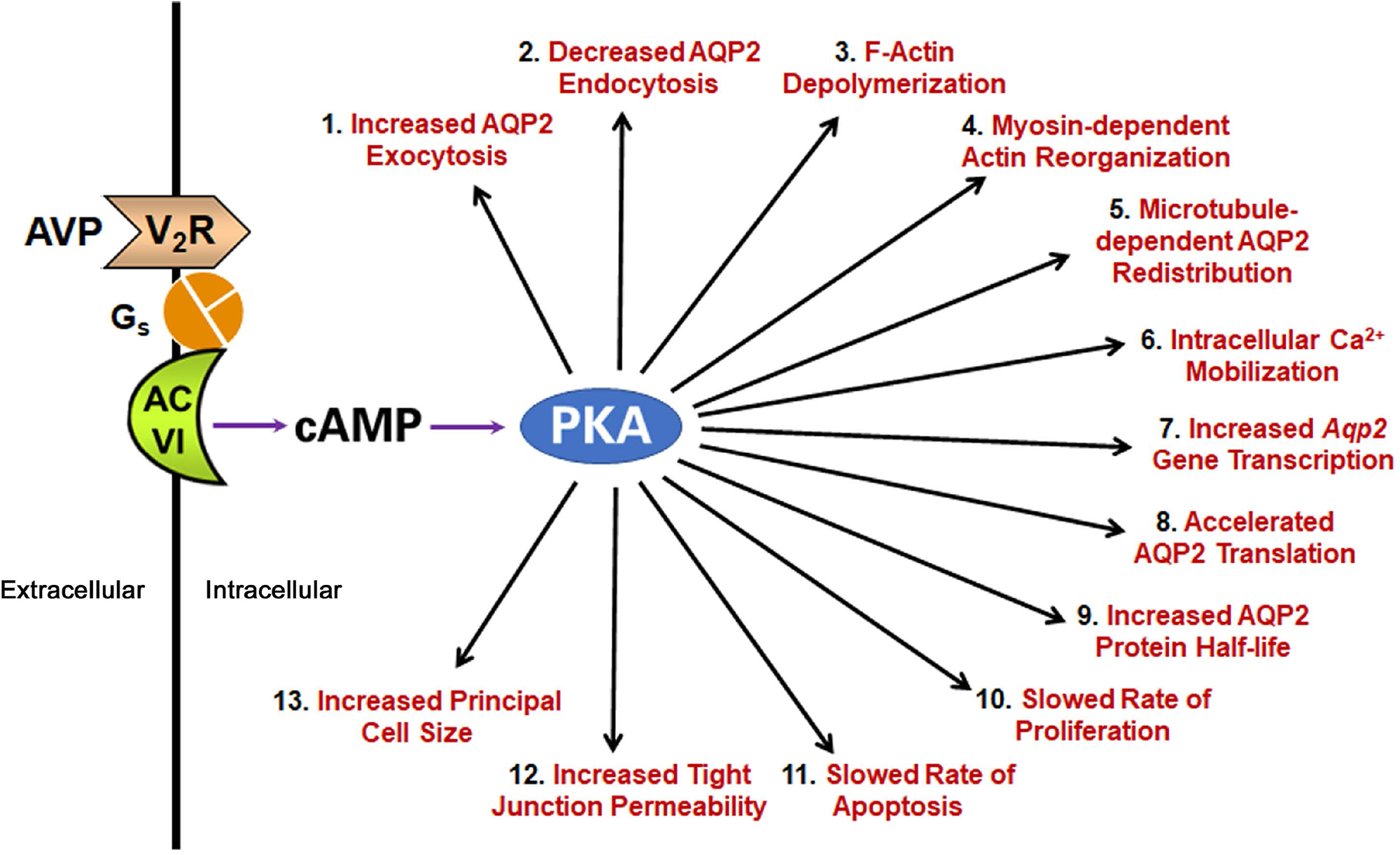
The physiologic responses at the cellular level elicited by vasopressin in the collecting duct epithelial cells as documented in prior literature (81).

## METHODS

The raw data used for this paper were reported as control experiments in our prior publication about PKA-independent signaling in collecting duct cells (35), but were not analyzed bioinformatically with respect to physiological mechanisms. For the reader’s convenience, we provide the methods used in a supplemental file (Supplemental Information – Methods). Briefly, mpkCCD, a collecting duct cell line, was grown on permeable supports using the Stable Isotope Labeling with Amino acids in Cell culture (SILAC) protocol for labeling (35). The cells were exposed to the vasopressin analog dDAVP (0.1 nM for 30 min) versus vehicle in three pairs of SILAC experiments. The two members of each pair, labeled with heavy and light amino acids, were mixed and subjected to our total and phosphoproteomic analysis protocol. Phosphopeptides were enriched using TiO_2_ and Fe-NTA columns. The enriched samples were analyzed using LC-MS/MS by HCD and EThcD fragmentation. The raw data was analyzed using Proteome Discoverer 1.4 (version 1.4.0.288). The peptide False Discovery Rate was set to <0.01(35).

### Phosphoproteomics Data Integration across Biological Replicates

For phosphoproteomics, the phosphopeptides having an area (MS1 scan) of at least 1.0E7 in each of the replicate cell clones along with a corresponding median ratio calculated from TiO_2_ and Fe-NTA results were considered for further analysis. The data files obtained from HCD and EThcD fragmentation were combined. For duplicate peptides, the ones with lowest standard deviation among three replicate clones were selected. Co-efficient of variation (CV) was calculated to provide an estimate of the biological variation among three independent clones. Phospho-peptides were considered changed in abundance if they met the following criteria:| Log_2_(dDAVP/vehicle) |>0.4 and (-log_10_(CV))>0.4 (35). Amino acid sequences (13-mer) of phosphopeptides were centralized around the phosphorylation site using PTM Centralizer (https://hpcwebapps.cit.nih.gov/ESBL/PtmCentralizer/).

### Generation of Sequence Logos

*PTM-Logo* (36) (https://hpcwebapps.cit.nih.gov/PTMLogo/) was used to generate sequence logos. The inputs were centralized sequences of specific curated groups of phosphorylation sites. Background sequences and chi-square alpha values for generation of each logo are noted in the figure legends.

### Bioinformatics Analysis

#### External Data Sources

Known substrates of various kinases and their regulated phosphorylation sites were downloaded from *PhosphoSitePlus* (https://www.phosphosite.org/homeAction). Known effects of phosphorylation changes were downloaded from *PhosphoSitePlus* (https://www.phosphosite.org/staticDownloads). Additional information about the effects of specific phosphorylation events was obtained from *Kinexus PhosphoNET* (http://www.phosphonet.ca/) and through direct *PubMed* searches. A list of PDZ domain-containing mouse proteins was downloaded from the *SMART* database (http://smart.embl.de/) (37). *Abdesigner* (https://esbl.nhlbi.nih.gov/AbDesigner/) was used to visualize the domain organization of proteins relative to the regulated phosphorylation sites (38).

#### Network Construction

STRING (https://string-db.org/) was used to create a relational network for proteins with increased and decreased phosphorylation sites using default settings to construct a preliminary signaling network. This network was the starting point for manual curation based on knowledge gleaned from prior literature to classify vasopressin signaling into various cellular processes, functions, or components. *Cytoscape 3.7.1* (https://cytoscape.org/) was used for visualization of the resulting networks.

#### Kinase Tree Annotation and Substrate Motif Generation

*KinMap* (39) (http://kinhub.org/kinmap/) was used to locate vasopressin-regulated protein kinases to the Manning dendrogram (40). Curated target sequences for each kinase (downloaded from *PhosphoSitePlus*) along with the kinases on the same sub-branch of the dendrogram were put into *PTM-Logo* to identify target sequence preferences for individual kinase neighborhoods. Background sequences for this analysis were the list of all phosphorylation sites in the *PhosphositePlus* mouse database. This analysis was visualized using *CORAL* (http://phanstiel-lab.med.unc.edu/CORAL/) (41). *CORAL* is an interactive web application for kinome annotation and visualization that is capable of generating vector graphics (SVG) images of the Manning kinome tree with customizable branch color and node color.

#### Gene Ontology (GO) Analysis

GO terms, annotated protein domains, and FASTA sequences were obtained using *ABE* (*Automated Bioinformatics Extractor*; http://helixweb.nih.gov/ESBL/ABE/). Chi-square analysis was used to identify the association between regulated proteins and various annotation terms that were obtained through *ABE*. InterPro database in the functional annotation tool, *DAVID* (https://david.ncifcrf.gov/) was used to determine the enrichment of specific protein domains. *DAVID* uses a modified Fisher Exact test (EASE score) to test the enrichment of annotation terms. The full list of unique accession numbers identified from the current phospho-proteomics experiment was used as a background for *DAVID* analysis.

#### Rule-based Classification of Regulated Phosphosites

All regulated phosphorylation sites were classified based on two criteria: 1) direction of change in phosphorylation following dDAVP stimulation; 2) the presence of specific amino acids in different positions upstream or downstream of altered phosphosites as described in the Results section.

### Data Availability

The raw files (.raw) can be accessed at www.ebi.ac.uk/pride/archive/ with the dataset identifier PXD015719. The curated data are available at https://hpcwebapps.cit.nih.gov/ESBL/Database/mpkCCD-AVP/.

## RESULTS

### General Observations

To identify vasopressin responses and regulated protein kinases in mouse renal mpkCCD epithelial cells, we used mass spectrometry (SILAC labeling) for proteome-wide quantification of phosphopeptides in three separate clones of the mpkCCD cells. The data were previously reported as control experiments for studies of mpkCCD cells in which PKA was deleted (35), but have not until now been presented explicitly or interpreted. The experiments described here compared phosphoproteome of PKA-intact mpkCCD cells with 30-minute exposure to the V2 receptor-selective vasopressin analog dDAVP (0.1 nM applied only to the basolateral side) to cells exposed only to vehicle. Prior time course studies established that 30 minutes is required for the steady-state response to vasopressin in collecting duct cells (14). The full dataset is provided to users at https://hpcwebapps.cit.nih.gov/ESBL/Database/mpkCCD-AVP/.

Figure 2A shows a torch plot indicating a general view of the phosphoproteomic responses following dDAVP stimulation. Of the 9640 quantified phosphopeptides (2649 phosphoproteins), 429 were classified as showing high probability abundance changes in response to dDAVP based on dual criteria (-Log_10_(CV)≥0.4 and |Log_2_(dDAVP/Vehicle) |>0.4) where CV means coefficient of variation (Supplemental Table 1). These phosphopeptides are found in 295 distinct phosphoproteins. Among the 429 phosphopeptides, 187 were increased and 242 were decreased according to the thresholds defined above. The 429 regulated phosphopeptides consisted of 332 mono-phosphopeptides and 97 multi-phosphorylated peptides. Among mono-phosphopeptides, 153 were increased and 179 were decreased.

**Figure 2.**
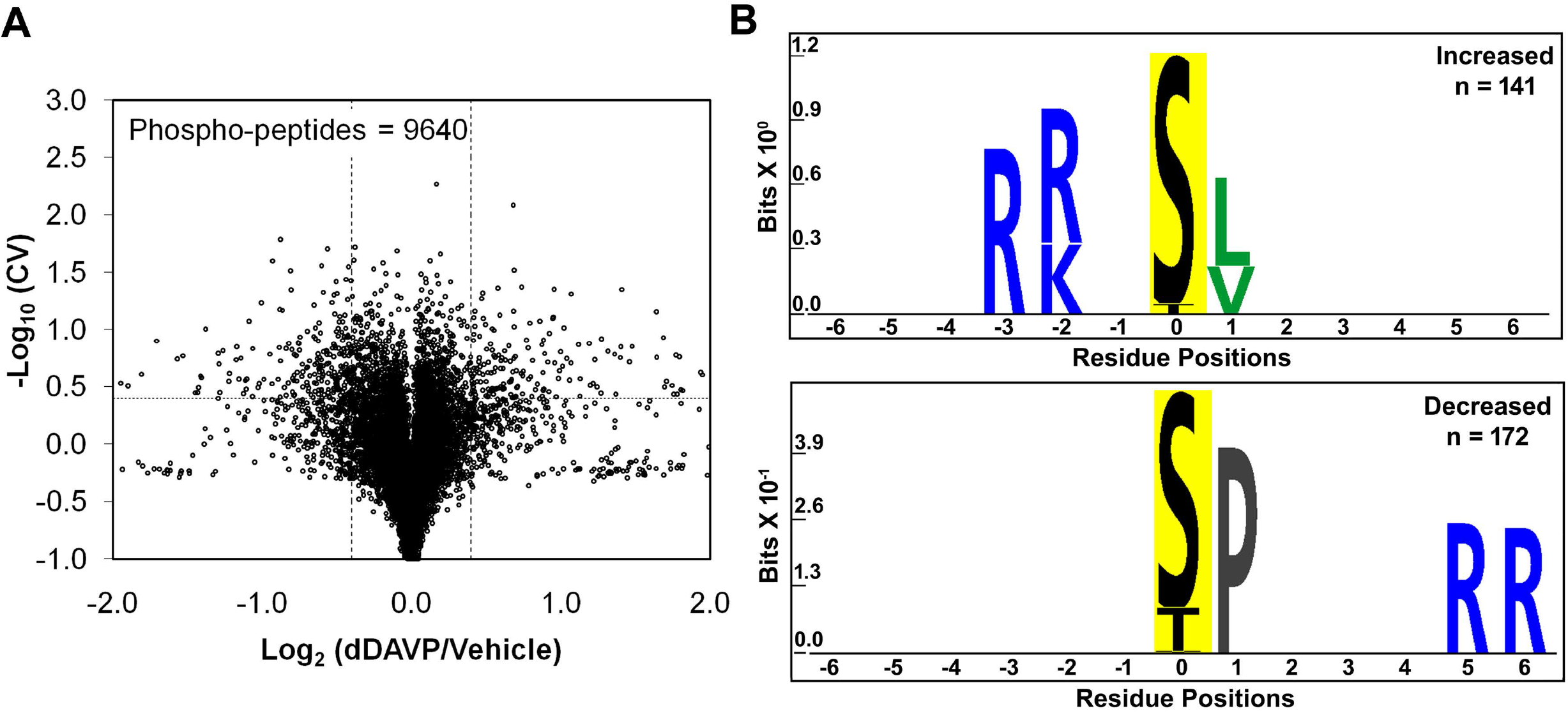
Vasopressin effects on phosphoproteome of mpkCCD cells. A. Torch plot for phosphopeptides quantified across all three pairs (dDAVP vs. vehicle) of mpkCCD cell clones. Vertical dashed lines show log_2_(dDAVP/Vehicle) of 0.4, and −0.4, while the horizontal line represents −log_10_(CV) of 0.4 (CV=0.4). B. Motif analysis shows the amino acid residues over-represented in sets of unique mono-phosphopeptides whose abundances are significantly increased (upper) and decreased (lower). *PTM-Logo* was used with whole mouse dataset as background and Chi-square cutoff was set at 0.0001.

In the same experiments, we also carried out total protein quantification for 10,128 proteins in mpkCCD cells, of which 7,339 proteins were quantified in all three pairs of samples (Supplementary Table 2). In general, treatment of the cells for 30 minutes with dDAVP resulted in few changes in protein abundances none of which impinge on phosphopeptide quantification.

To identify general sequence preferences among phosphorylation sites that respond to vasopressin, we used *PTM-Logo* (36) inputting the 13-mer sequences surrounding the 313 unique phosphosites in the 332 altered mono-phosphopeptides from Figure 2A (Figure 2B). The increased phosphorylation sites (n=141, corresponds to 153 mono-phosphopeptides) identified a motif typical of basophilic protein kinases (members of AGC and CAMK families, namely [R-(R/K)-X-*p*(S/T)] (Figure 2B, upper panel). This motif is commonly viewed as a PKA target motif, but can be seen in targets of other basophilic kinases (42). The logo generated from the decreased sites (n=172, corresponds to 179 mono-phosphopeptides) in the mpkCCD cells identified a preference for a proline in position +1 relative to the phosphorylated amino acid (Figure 2B, lower panel). This is consistent with reduced activity of one or more proline-directed kinases from the CMGC family, which include mitogen-activated protein kinases and cyclin-dependent protein kinases (42).

We mapped all altered phosphoproteins to specific cellular functions using STRING and GO term annotations (Figure 3). The increased phosphopeptides are indicated as green nodes and the decreased ones are indicated as red nodes. A vector-graphic, zoomable version of this figure can be accessed at https://hpcwebapps.cit.nih.gov/ESBL/Database/Vasopressin-Network/. Many of the functional groups defined in this way correspond closely to known cellular level actions of vasopressin in collecting duct cells (Figure 1). Some categories of proteins define other possible roles of vasopressin not previously identified in the literature, e.g., “Desmosome”, “mRNA Stability”, “mRNA Processing”, and “Nuclear Envelope”. A particularly large number of phosphoproteins are involved in “Gene Transcription”.

**Figure 3.**
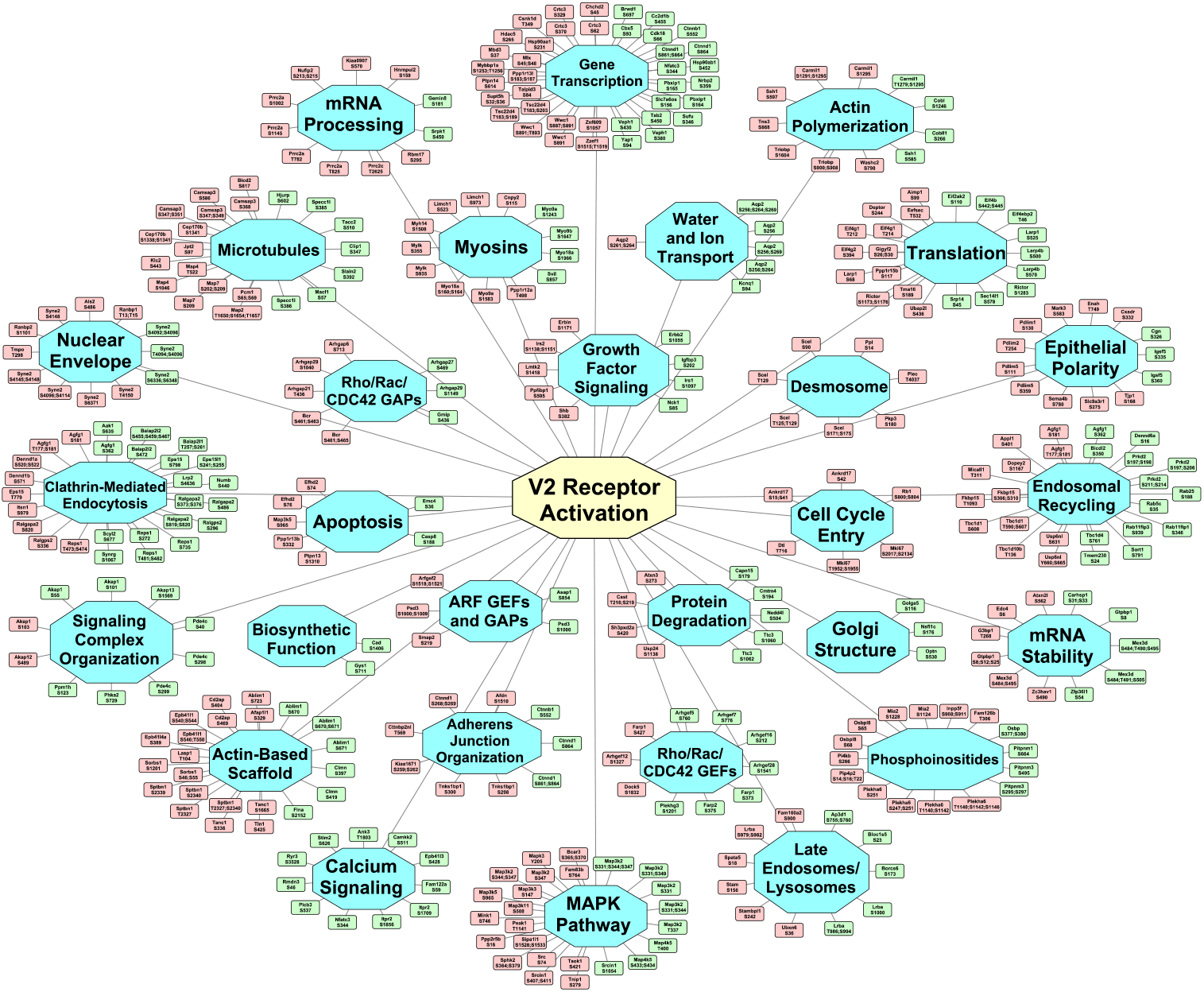
Vasopressin signaling network. Undirected graph depicting the mapping of proteins containing altered phosphorylation sites to specific cellular functions or structures [blue nodes]. The green and red nodes indicate specific phosphorylation sites altered in response to dDAVP administration (green, increased; red, decreased). A vector-graphic, zoomable version is provided at https://hpcwebapps.cit.nih.gov/ESBL/Database/Vasopressin-Network/ to aid in viewing details. Network was constructed initially in STRING and manually extended to the full data set using GO terms and *PubMed* entries. *Cytoscape* was used for visualization.

### Vasopressin-responsive Protein Kinases

Of the 295 vasopressin-regulated phosphoproteins, 25 are themselves protein kinases that are candidate kinases for downstream vasopressin signaling. These are listed in Table 1. These kinases are mapped to a *KinMap*-based dendrogram (Figure 4), classifying kinases on the basis of similarities of their catalytic regions. Different branches define kinase families that typically share target sequence preferences. Interestingly, none of the kinases that undergo phosphorylation changes are in the AGC category, but instead are chiefly members of the CAMK, CMGC, TLK, CK1, STE, and TK families as well as a few that are unclassified. The kinases shown in Table 1 are, of course, not the only kinases that are regulated in mpkCCD cells because differential phosphorylation is but one of many modes of regulation. Several of the sites that changed have known effects on the catalytic activity, namely Mapk3 at Y205, Src at S74, Lmtk2 at S1418, Camkk2 at S511 and Mylk at S355 and S935. Most of the remaining sites either have not been directly studied with regard to effect on kinase activity or the effects have not been reported on *PhosphositePlus* or *PhosphoNET*. Of special note is ERK1 (Mapk3), which undergoes a decrease in phosphorylation at its active site, indicating decreased activity of ERK1 in response to vasopressin. ERK1 is a proline-directed kinase and a decrease in its activity provides a likely explanation for the X-*p*(S/T)-P motif prevalent among decreased phosphorylation sites (Figure 2B). Also, of interest are phosphorylation changes in multiple MAP kinase kinase kinases (Map3k2, Mapk3k3, Map3k5 and Map3k11) which can be upstream from MAP kinase activation (p38, JNK and ERK proteins).

**Figure 4.**
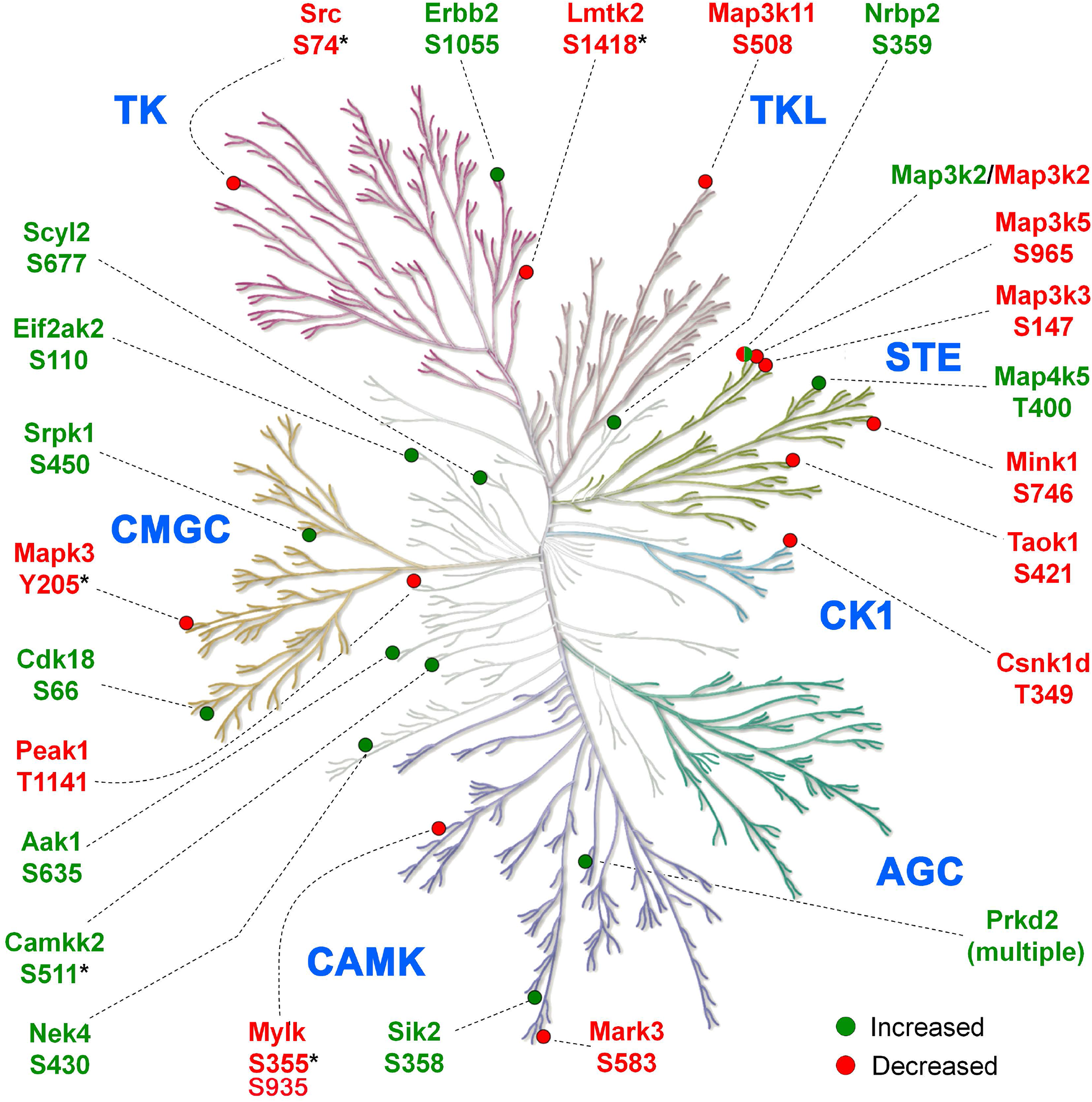
Vasopressin responsive phosphorylation sites mapped to protein kinases. Green and red bullets indicate increased and decreased phosphorylation sites respectively following dDAVP treatment. *KinMap* was used to generate the kinome tree. *, phosphorylation sites with known effects on catalytic activity.

**Table 1.**
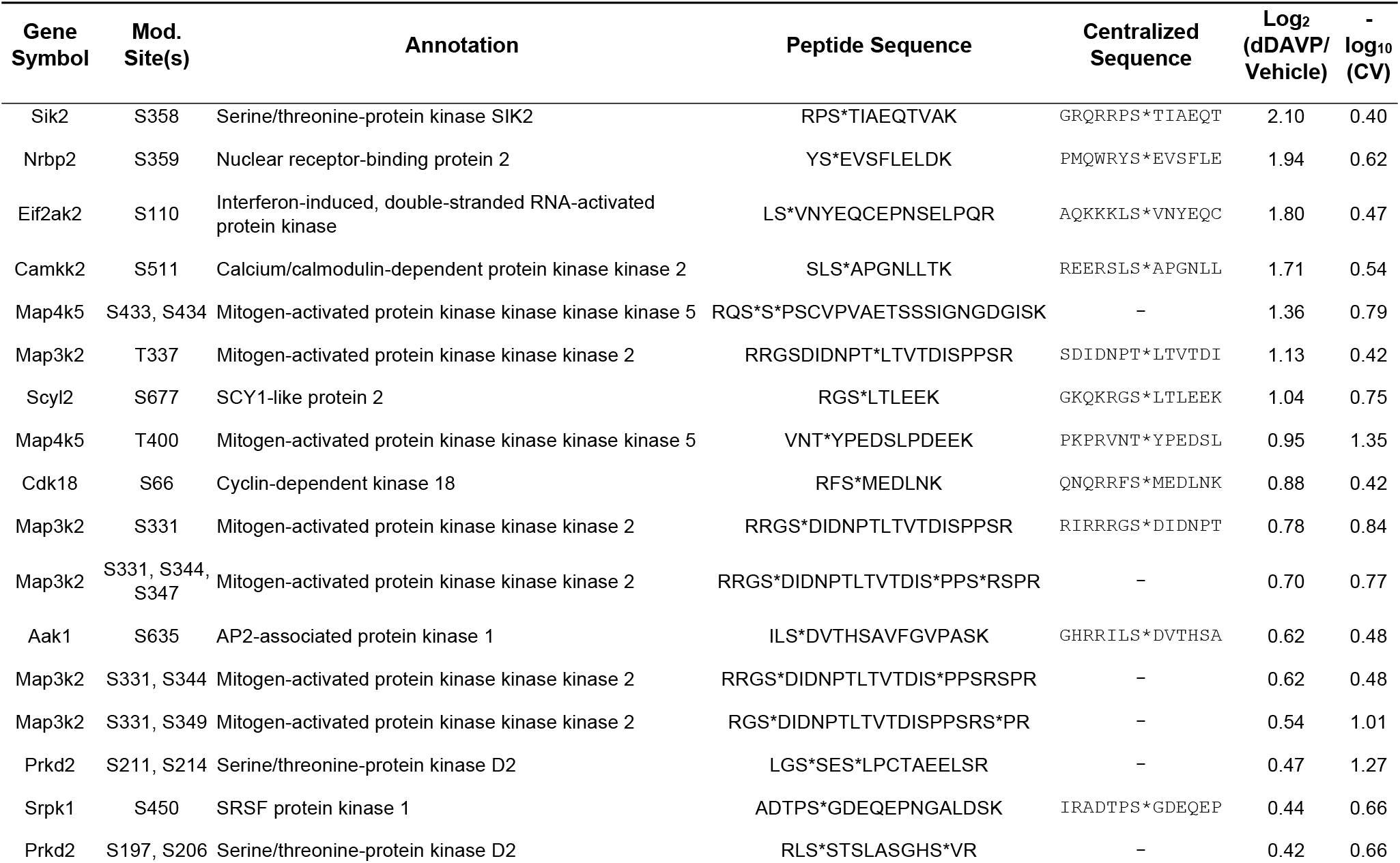
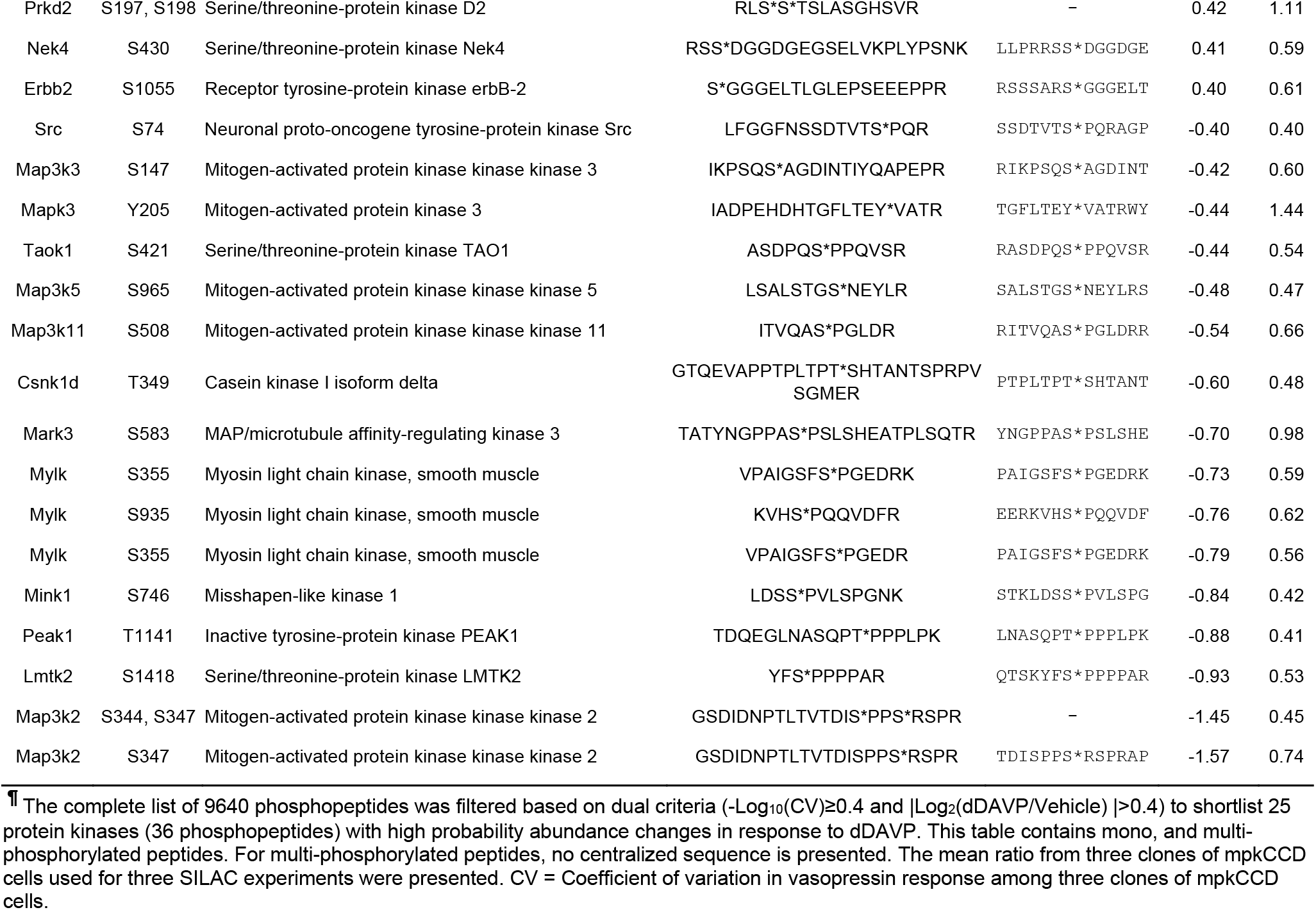
Protein kinases with vasopressin-responsive phosphopeptides in mpkCCD renal epithelial cells^¶^.

### Target Motifs of Vasopressin-responsive Protein Kinases

To predict target motifs for each regulated kinase we used the sequences of known targets of all kinases (data downloaded from *PhosphoSitePlus*) on a particular branch of the kinase dendrogram in Figure 4. For example, since Sik2 and Mark3 (SNF1-subfamily kinases) underwent a change in phosphorylation, we collected the centralized sequences of known targets of Mark3, Sik2, plus 11 other kinases (274 sequences, Supplemental Table 3) from the same subfamily (same branch of dendrogram) and used *PTM-Logo* to create a sequence preference prediction for Mark3/Sik2-related kinase. The resulting logos along with the branches color-coded to indicate the involved protein kinases are shown on Figure 5. The list of kinases for various sub-groups, total number and the list of centralized sequences of substrates for each of these groups that were retrieved from the *PhosphoSitePlus* database are provided in Supplemental Table 3. Of note, several vasopressin-regulated kinases were excluded from this analysis due to an insufficient number (i.e. less than 25) of known substrates, viz., Nrbp2, Camkk2, Aak1, Lmtk2, Peak1 and Scyl2. Conversely, PKA was included due to its central role in the V2R signaling although we did not detect altered phosphorylation of either of the two catalytic subunits (Prkaca and Prkacb) in the current study. This analysis offered a few interesting results. E.g. 1) Despite being a member of the CAMK family, the substrates of Protein Kinase D (Prkd2)-related kinases share a motif [L-X-R-(R/H)-X-*p*(S/T)] that resembles the PKA target motif. 2) Not all members of CMGC family of kinases are proline-directed. The substrates of Srpk1-related kinases (Srpk1-Srpk2, Clk1-Clk4) tend to have an arginine (R) in position −3 and −1, as seen previously using *in vitro* kinase assay of dephosphorylated HeLa cell lysate (43).

**Figure 5.**
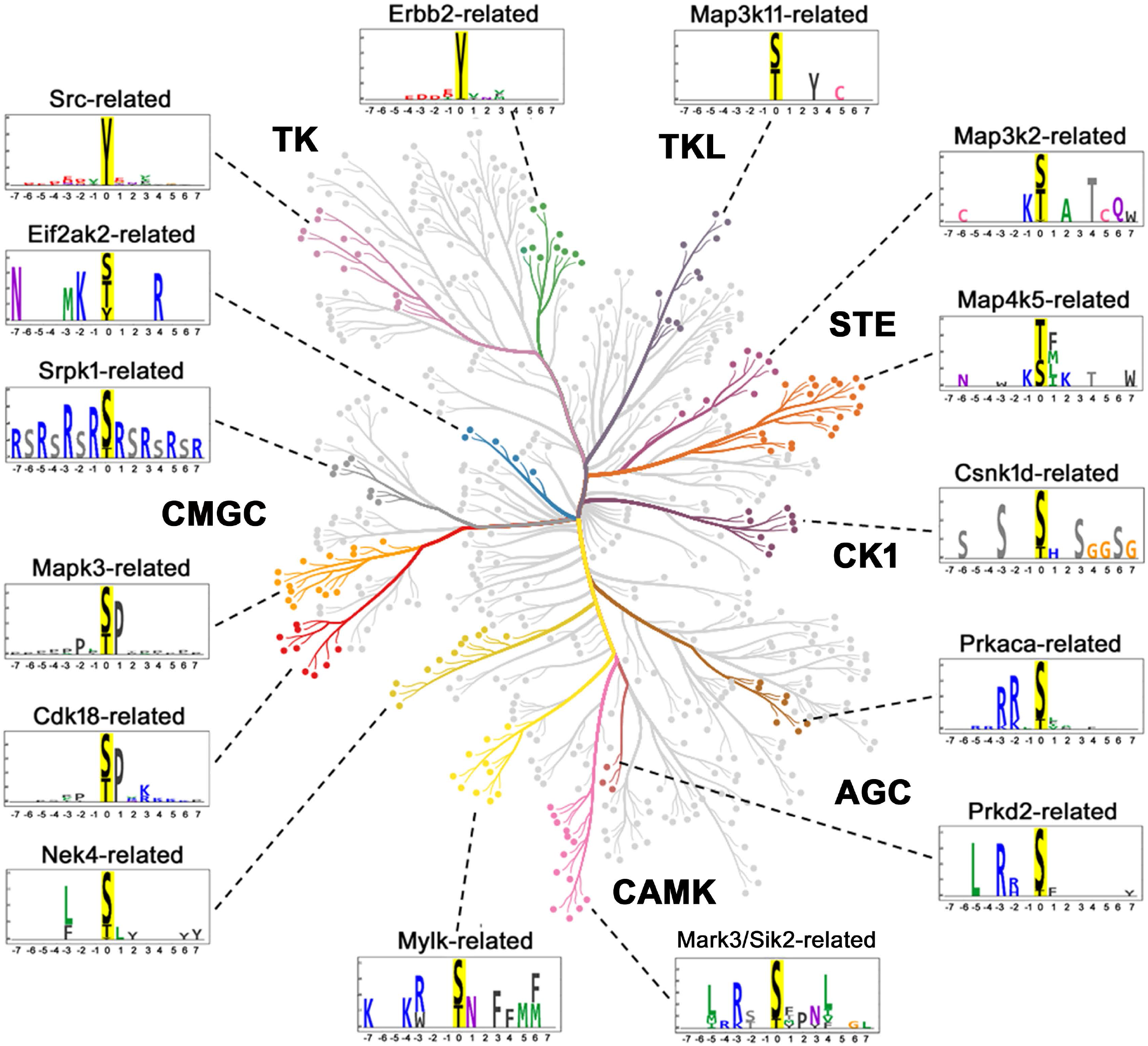
Substrate motifs corresponding to different branches of kinase dendrogram (color-coded) that contains one or more vasopressin responsive protein kinase as depicted in Figure 4. Prkaca (PKA) being the central kinase in V2R signaling was added although altered phosphorylation of PKA was not detected in the current study. The logos were generated after combining known substrates of all kinases in a particular branch that are available in the *PhosphoSitePlus* database. *CORAL* was used to generate the color-coded kinome tree. *PTM-Logo* was used for logo generation with 15-mer centralized peptide sequences. Whole mouse proteome was used as background for motif analysis. Chi-square = 0.001

### Rule-based Classification of Regulated Phosphorylation Sites

Figure 6 (top panel) presents a rule-based classification of the 313 regulated phosphorylation sites based on the sequence surrounding the phosphorylated S or T and the direction of change, yielding 11 subgroups (Supplemental Table 4). The over-representation of specific amino acids (arginine (R), lysine (K), and proline (P)) in the motif analysis (Figure 2B) guided this classification. One objective was to assign one or more protein kinases from Figure 4 to individual regulated phosphorylation sites. The substrate logos generated in Figure 5 for vasopressin-regulated kinase sub-groups were used to find the candidate upstream kinases. The candidate kinases for these sub-groups are listed in Figure 6 (bottom panel). A total of 133 out of the 313 phosphosites possessed a P in position +1. A majority of these sites (91%, n=121, sub-group 1) displayed reduced phosphorylation following dDAVP treatment (predicted kinase: Mapk3). However, the presence of at least 12 of these sites with increased phosphorylation in response to vasopressin (sub-group 2) indicates likely activation of one or more proline-directed kinase(s) from the CMGC family. The predicted kinase for these targets is Cdk18. The phosphosites without P at position +1 were next divided into two groups; phosphosites with (n= 110) and without (n=70) positively charged amino acids, R, K, and histidine (H) in position −3. This step was done to separate substrates of nominally basophilic kinases from non-basophilic kinases. Out of these 110 phosphosites, 93 had increased phosphorylation (85%), while 17 (sub-group 3) had decreased phosphorylation (15%) in response to vasopressin. The 93 increased sites were classified based on the presence of R/K/H in position −2. Among these, 63 (68%, sub-group 4) had R/K/H in position −2. These are likely target sites of PKA or possibly protein kinase D (Prkd2). Interestingly, in more than half (16 out of 30, sub-group 5) of the sites that do not have R/K/H in position −2, R/K is present in position −5, pointing to possible roles of Sgk1, Akt1 or Lats1 (44) although none of these kinases are reported in Table 1 as undergoing significant changes in phosphorylation in response to vasopressin. Among the upregulated 36 sites that do not contain R/K/H in position −3, two major sub-groups can be identified based on whether R/K is present in position −2. Sixteen sites have R/K in position −2 (44%, sub-group 7, no predicted kinase). Out of the remaining 20 sites that do not have R/K in position −2, 15 had S/D/E in position −3 (no predicted kinase, sub-group 8). Subgroup 10 contains 12 out of 34 (33%) down-regulated sites that contain S/T in position −3 instead of R/K/H (predicted kinase: Csnk1d). Overall, about 81% (Figure 6, 58/313) of the phosphorylation sites were mapped to candidate upstream kinases. The remaining 19% of the substrates with no predicted kinase may be targeted by kinases predominantly on the basis of co-localization.

**Figure 6.**
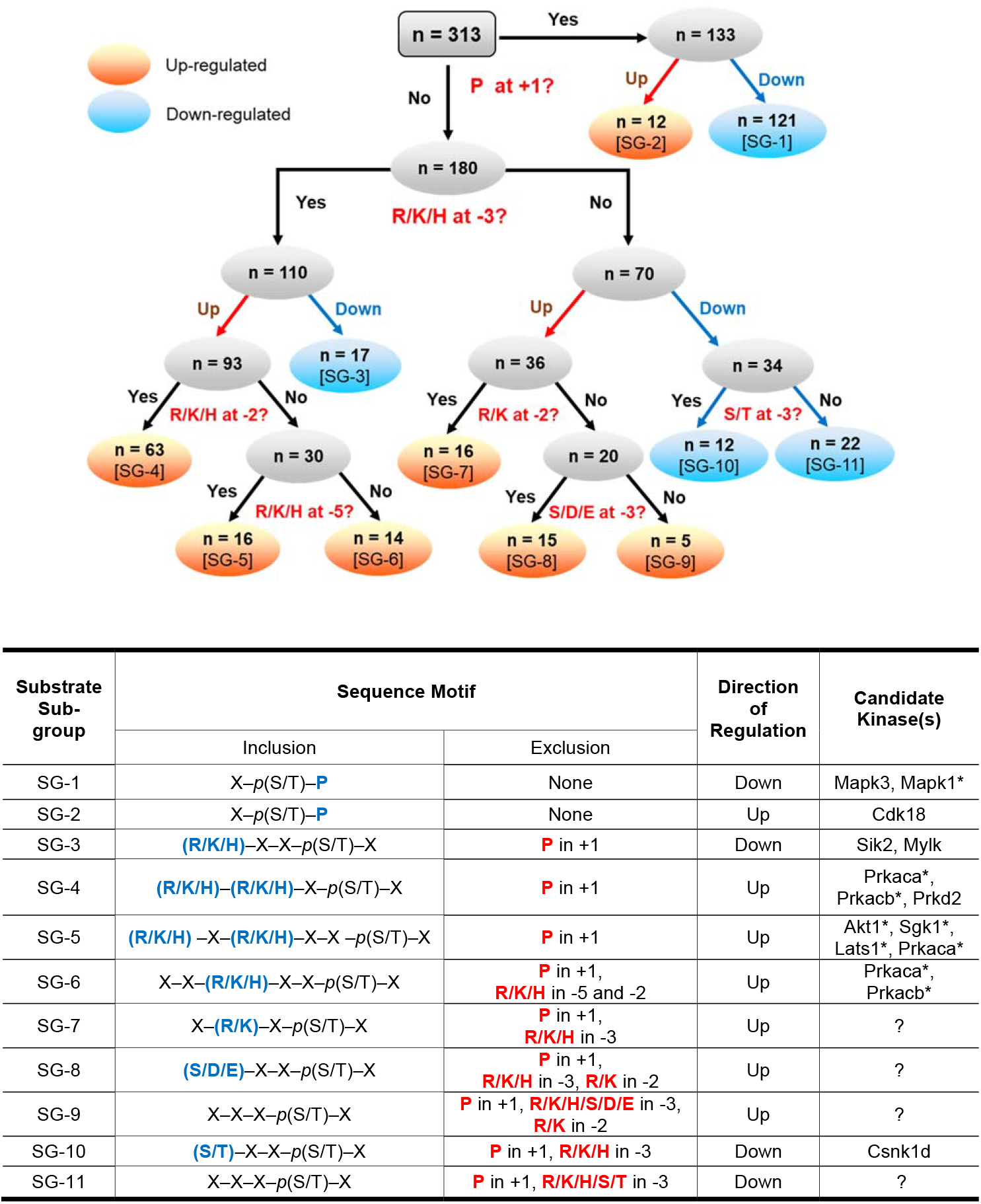
Rule-based classification of vasopressin-regulated phosphorylation sites. **Top panel:** All 313 regulated single phosphorylation sites were classified into 11 sub-groups (numbered SG-1 to SG-11 in square brackets). This classification is based on the direction of change (up-regulated or down-regulated) and sequence surrounding the phosphorylated S or T in position +1, −2, −3 and −5. Seven sub-groups contain up-regulated phospho-sites (orange) while 4 subgroups contain down-regulated phospho-sites (blue). **Bottom panel:** The table presents the sequence motifs in the ‘inclusion’ column (amino acids in blue color) for all 11 sub-groups of substrates along with the candidate kinases. Amino acid residues that are excluded from specific positions are indicated in the exclusion column. *Kinases that are not listed Table 1. ‘?’, no known kinase.

### Co-localization of Kinase and Target Substrates

Intuitively, for a phosphorylation reaction to take place, a protein kinase and its substrate must be in the same sub-cellular compartment. Figure 7 shows a heat map representation showing relative abundances of the regulated kinases from Table 1 in subcellular fractions from mpkCCD cells. The data for regulated substrates is reported in Supplemental Table 5. These data were previously reported as a web resource in a study in which we employed differential centrifugation (1K, 4K, 17K, 200K-pellet and 200K-supernatant) and protein mass spectrometry to estimate subcellular distribution of individual proteins of mpkCCD cells following short-term V2R stimulation (45, 46). As can be seen in Figure 7, some kinases are highly restricted regarding subcellular localization. For example, Camkk2, which is present mainly in the cytosolic fraction may not be a good candidate for phosphorylation of AQP2, which is absent from the cytosolic fraction.

**Figure 7.**
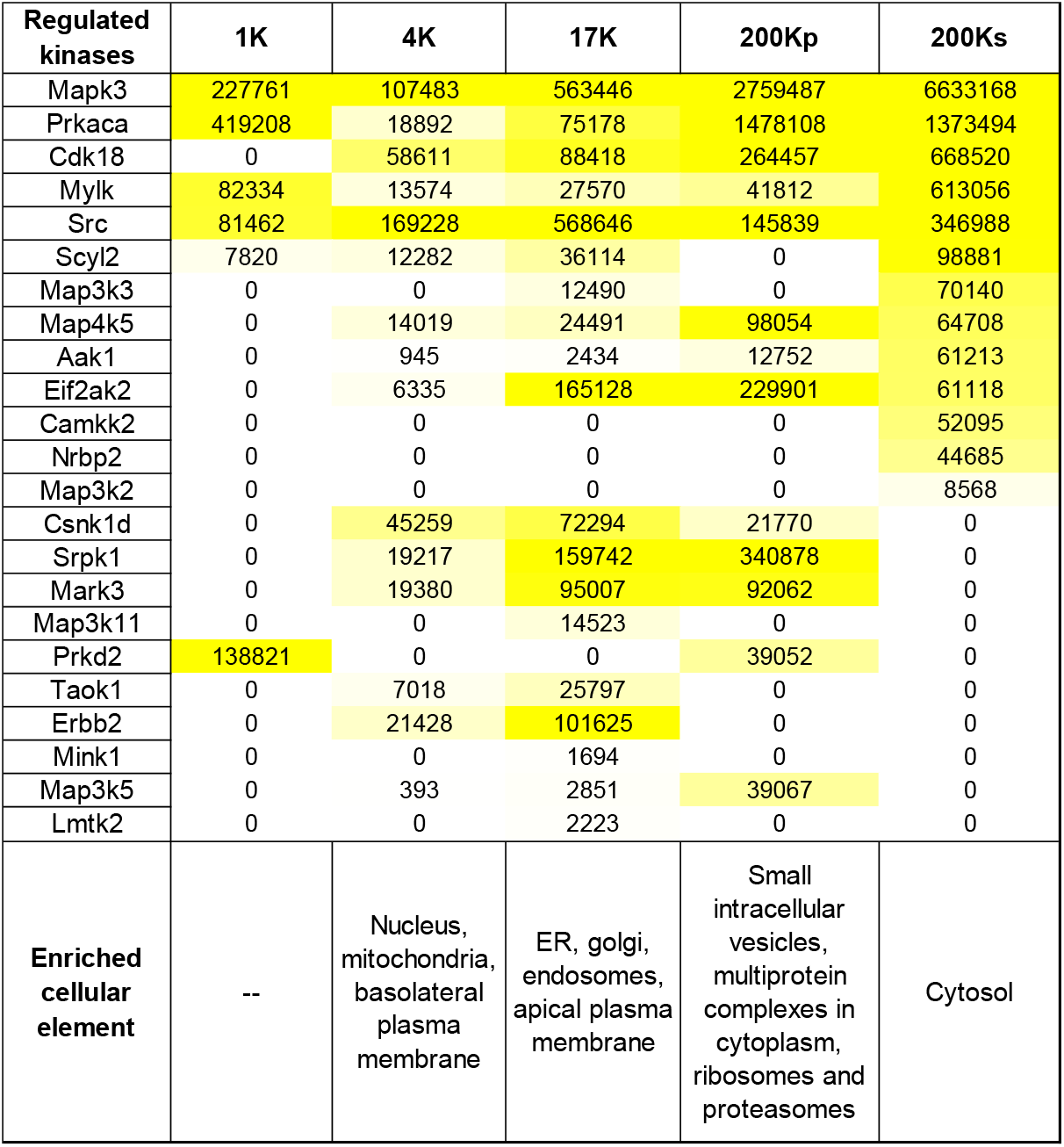
Protein abundances of altered kinases in differentially centrifuged fractions from mouse mpkCCD cells after treatment with 0.1 nM dDAVP for 30 min. Abundances were calculated from total areas under the reconstructed chromatograms. The area was normalized with the molecular weight of the respective protein kinase and presented in arbitrary units. The values were color-coded to relative abundance level to aid visualization. The data was obtained from a web resource generated by Yang et al. (45, 46). These fractions were loosely assigned to various elements of the cell based on prior literature evidence and distribution of the marker proteins in the above two studies.1K, 1000 Xg; 4K, 4000 Xg; 17K, 17,000 Xg; 200Kp, 200,000 Xg-pellet; 200Ks, 200,000 Xg-supernatant.

### Subcellular Targeting Mechanisms for Kinases and their Substrates

Some protein kinases such as PKC isoforms are preferentially active at membrane surfaces (47). We identified “membrane-associated proteins” as those with *GO Cellular Component* terms containing the strings “membrane”, “vesicle” or “endosome”. 93 of 137 phosphoproteins with increased phosphorylation were membrane-associated proteins (68%) compared with 1163/2512 (46%) of all phosphoproteins that did not contain upregulated phosphopeptides (Chi-square, P=0.000), consistent with preferential phosphorylation at membrane surfaces (see Supplemental Spreadsheet 1 for protein lists). Similar findings were obtained with phosphoproteins that contain phosphopeptides decreased in response to dDAVP, in which 109 proteins out of 183 (60%) were membrane-associated (Chi-square, P=0.001, Supplemental Spreadsheet 1).

To identify possible mechanisms of membrane interaction, we used *DAVID* to identify proteins with lipid-binding or protein-binding domains that are enriched in upregulated phosphopeptides. This analysis identified the pleckstrin homology (PH)-like domain as a predominant domain, (found in 22 of the 137 upregulated phosphoproteins, EASE=0.000; 28 out of 183 downregulated phosphoproteins, EASE= 0.000). PH domains mediate membrane binding of proteins through interactions with inositol phospholipids (48), but can also mediate binding to certain proteins such as protein kinase C (49). Prkd2 was found to be a PH domain containing kinase identified as vasopressin-regulated in the current study (Figure 8).

**Figure 8.**
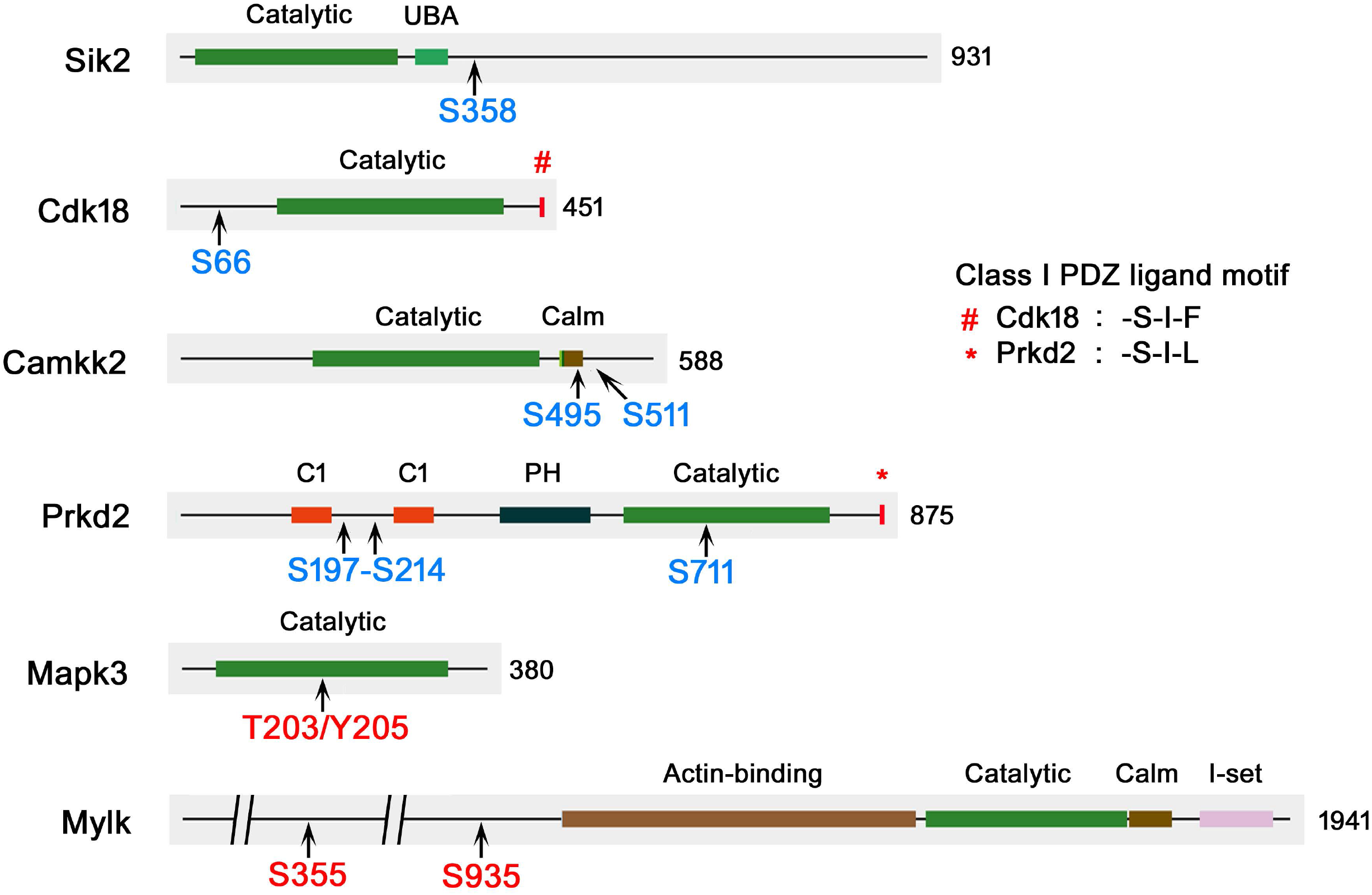
Protein kinases that undergo phosphorylation downstream from PKA. Locations of vasopressin-responsive phosphorylation sites are indicated in blue (increased) or red (decreased). Class I PDZ ligand motif on the carboxyl terminal is shown. The total number of amino acid residues in each kinase are shown. Calm, calmodulin-binding domain; C1, diacylglycerol-binding domain; PH, plekstrin homology phosphoinositide binding domain; UBA, ubiquitin-associated domain; I-set, Immunoglobulin I-set domain. *Abdesigner* (38) was used for domain visualization.

Another means of membrane targeting of proteins is interaction between PDZ domain-containing proteins and proteins with COOH-terminal PDZ ligand motifs, which are often integral membrane proteins (50). It is noteworthy that a Class I COOH-terminal motif is present in PKA catalytic subunit alpha (-T-E-F). Consequently, we analyzed the COOH-terminal sequences of upregulated and membrane-associated phosphoproteins to explore possible roles of PDZ interactions in membrane localization. The class I PDZ motif was defined as -(S/T)-X-Ø where S/T indicates serine or threonine, X indicates any amino acid and Ø indicates any hydrophobic amino acid (W/F/L/I/V/A/M) (51). A total of 13 out of 93 (14%) membrane-associated phosphoproteins with upregulated phospho-sites possessed COOH-terminal Class I PDZ ligand motifs compared with 91 of 1158 (8%) of non-upregulated membrane-associated phosphoproteins (Chi-square, P=0.040, Supplemental Spreadsheet 2). Among the vasopressin-regulated kinases in Table 1, Cdk18 (-S-I-F) and Prkd2 (-S-I-L) were found to possess Class I PDZ ligand motif on their carboxyl terminus (Figure 8). In addition, nine PDZ domain proteins underwent changes in phosphorylation namely, Afadin (Afdn), ZO-1 (Tjp1), Deptor, Pdlim1, Pdlim2, Pdlim5, Ptpn13, Sipa1l1 and NHERF1 (Slc9a3r1). Remarkably, all exhibited substantial decreases in phosphorylation and 8 out of 9 of these decreases were at sites with P in position +1 relative to the phosphorylated amino acid, consistent with a potential role of ERK.

A related question is the role of AKAP proteins in compartmentalization of protein kinase activities. Six different phosphorylated Akap proteins were identified, namely Akap1, Akap2, Akap8, Akap9, Akap12 and Akap13. Among these, Akap1, Akap12 and Akap13 underwent changes in phosphorylation in response to dDAVP (Figure 3). Two upregulated sites in Akap1 (S55 and S101) were increased at classical PKA target sites [R-R-X-*p*(S/T)]. AKAPs not only bind to PKA regulatory subunits but also other protein kinases, protein phosphatases, cyclic AMP phosphodiesterases, adenylyl cyclases, phospholipases and other signaling proteins (52). Among these, the analysis detected vasopressin-induced changes in phosphorylation of cyclic nucleotide phosphodiesterase Pde4c, protein phosphatase Ppm1h, protein phosphatase slingshot 1 (Ssh1), phospholipase C beta (Plcb3), protein kinase Src, Prkd2, and multiple protein phosphatase regulatory subunits. Several of these changes were in basophilic sites compatible with phosphorylation by PKA. For example, Pde4c underwent increases in phosphorylation at two putative PKA target sites, S40 and S299. Akap13 may also organize MAP kinase signaling through the Ksr1 anchor protein (53).

## DISCUSSION

Vasopressin exerts its physiological effects in collecting duct cells through a GPCR that signals largely through G_α_s-mediated activation of adenylyl cyclase 6 (3), production of cAMP and activation of PKA. Indeed, we previously showed that almost all vasopressin-mediated phosphorylation changes in mpkCCD cells are ablated when the two PKA catalytic genes (*Prkaca* and *Prkacb*) are deleted via CRISPR-Cas9 (PKA-null cell) (34). In the vasopressin signaling network, a majority of downstream actions can result from PKA-mediated regulation of other protein kinases from various kinase subfamilies that have different target sequence specificities and different localizations in the cell. Based on the direction of phosphorylation change and the sequence surrounding the phosphorylated amino acid, all 313 phosphorylation sites were classified into sub-groups that could be mapped to the regulated kinases (Figure 6). About 81% of phosphorylation changes are potentially mediated by just a few protein kinases, viz. Mapk3 (ERK1), PCTAIRE-3/Cdk18, Camkk2, Prkd2, Mylk and Sik2 in addition to PKA. Figure 8 shows the location of phosphorylation sites relative to the major protein domains in each of these six downstream kinases. Based on prior literature, these kinases appear to be the best candidates for further investigation of vasopressin signaling. Here we discuss these kinases with regard to prior literature both in collecting duct cells and in other cell types. This list is based on phosphorylation changes alone and several other kinases are known to be regulated by other mechanisms, e.g. activation by calcium mobilization through calmodulin such as calcium/calmodulin-dependent protein kinase 2δ (Camk2d) (54).

### Salt-inducible Kinase 2 (Sik2)

In our prior study in PKA-null cells (34), two sites in Sik2 showed striking decreases in phosphorylation, namely S358 and S587, both of them with sequences compatible with direct PKA phosphorylation. In this study, only S358 showed a significant increase in response to vasopressin (more than 4-fold), presumably PKA-mediated. PKA was previously demonstrated to phosphorylate S358 of Sik2 in adipocytes (55), reducing its enzymatic activity (56) via 14-3-3 binding (57). Thus, Sik2 activity is likely reduced by vasopressin in collecting duct cells. Sik2 is an AMPK-like kinase with a basophilic target motif as illustrated in Figure 5. Among its targets are CREB-related transcriptional coregulators Crebbp, Ep300, Crtc1, Crtc2 and Crtc3. All three of the CRTC proteins showed large increases in phosphorylation at Sik2 sites in response to deletion of both PKA catalytic genes (34). In this study, Crtc3 showed a marked decrease in phosphorylation at three recognized Sik2 sites demonstrated in prior studies, viz. S62, S329 and S370 (58). The CRTCs activate transcription of cAMP-responsive target genes by interacting with one or more of the three cAMP response element-binding protein (CREB) transcription factors (ATF1, CREB1 or CREM) (57). Therefore, the PKA-Sik2-Crtc3 regulatory pathway has potential relevance to the regulation of *Aqp2* gene transcription by vasopressin.

### Cyclin-dependent Kinase 18

Cdk18 is a proline-directed kinase (CMGC Family) (Figure 4) that we have identified in previous studies as a target for regulation by vasopressin (32, 59). The regulation is of two types. First, vasopressin treatment markedly accelerates transcription of the Cdk18 gene (32). This effect is likely PKA-dependent because Cdk18 protein abundance was markedly decreased in PKA-null cells (34). In addition, in native rat inner medullary collecting duct (IMCD) cells, short term exposure to vasopressin resulted in a marked increase in phosphorylation at S66 (60) at a site compatible with phosphorylation by PKA (Figure 8). In the present study in mouse mpkCCD cells, vasopressin increased S66 phosphorylation as well, giving further support to the conclusion that Cdk18 could be an important downstream target of PKA. Vasopressin did not increase phosphorylation at this site in PKA-null mpkCCD cells (35). In HEK293T cells, PKA mediated phosphorylation has been shown to activate Cdk18, with phosphorylation at three sites including S66 (61). If PKA does this in collecting duct cells, Cdk18 activation would provide a likely explanation for the many phosphorylation sites with proline in position +1 that increase in response to vasopressin (Figure 6). Cdk18 expression in collecting duct has recently been confirmed by Dema et al. (62) who proposed a role in regulation of AQP2 degradation. We found that Cdk18 is very abundant in nuclear fractions in both mpkCCD (63) and native IMCD cells (64) and speculate that it could be involved in regulation of transcriptional events in collecting duct cells as well.

### Calcium/calmodulin-dependent Protein Kinase Kinase 2

Camkk2 is a calcium/calmodulin-dependent protein kinase (Figure 8) known to phosphorylate various downstream targets including AMP-activated kinase (65). Our early phosphoproteomics findings in suspensions of native rat IMCD cells identified Camkk2 as a component of vasopressin signaling (66). Those studies demonstrated that vasopressin increases phosphorylation at two sites, S494 and S510 (equivalent to S495 and S511 in mouse). Studies in PKA-null cells showed decreases in phosphorylation at both sites, with almost total ablation of phosphorylation at S495 (34). This confirmed the conclusion from studies in COS-7 cells that both S495 and S511 phosphorylation are PKA-dependent (67). The phosphoproteomic results in the current study of PKA-intact mpkCCD cells are consistent with the prior findings, providing evidence for vasopressin-mediated phosphorylation at both sites with the greatest increase in S511 phosphorylation. Vasopressin potentially has two effects on Camkk2 activity in the collecting duct: 1) activation via vasopressin-induced calcium mobilization (demonstrated in native collecting duct) (22); and 2) inhibition via S511 phosphorylation, which works by interfering with calcium/calmodulin binding to an activation site (68) (Figure 8). Camkk2 is found in nuclear fractions in mpkCCD cells (63) and native rat IMCD cells (64) and therefore is likely to phosphorylate nuclear proteins. For example, Camkk2 has been found to be required for activation of the transcription factor cyclic AMP-responsive element binding protein (CREB1) in the hippocampus (69) and if true in collecting duct cells such a response could be involved in vasopressin-mediated regulation of *Aqp2* gene transcription.

### Protein Kinase D2 (Prkd2)

Prkd2 is a diacylglycerol regulated kinase that underwent increases in phosphorylation at several sites (Figure 8). Previously, in PKA-null cells, we found markedly decreased phosphorylation at S197 (34), suggesting that it could be a direct PKA target site (centralized sequence: ARKRRLS*STSLAS). Here, we found increases in two phospho-S197-containing double phospho-peptides. Also in prior studies in native rat IMCD cells, we found a moderate increase at another site, S711 (59), which nominally matches a moderate increase in the present experiments. Phosphorylation of this site has been reported to activate Prkd2 (70). Thus, although the evidence is not ironclad, it suggests that vasopressin increases Prkd2 activity. The Prkd2 target motif is similar to that of PKA (Figure 5), Thus, some increases in phosphorylation attributed to PKA could also be due to Prkd2 activation.

### Mitogen-activated Protein Kinase 3 (Mapk3)

ERK1 (Mapk3) is a serine/threonine kinase which along with ERK2 (Mapk1) plays an important role as the endpoint of the classic Raf-MEK-ERK cascade. ERK stimulates cell proliferation and dedifferentiation through phosphorylation of AP1 and Ets family transcription factors (71, 72). It also has many effects on cytoplasmic processes. ERK1 is regulated by active site phosphorylation. In this study, a moderate decrease was observed in active site phosphorylation of Mapk3, with a smaller response in Mapk1. This confirms earlier observations of decreases in active site phosphorylation in response to vasopressin in cultured MDCK cells (73), rat IMCD (74) and mpkCCD cells (75). These responses can help to explain the anti-proliferative, pro-differentiation effects of vasopressin in normal collecting duct cells (76). In PKA-null cells, active site phosphorylation was increased compared to PKA-intact mpkCCD cells indicating that the decreases seen with vasopressin are likely downstream from PKA activation (34). ERK1 and ERK2 are proline-directed kinases with a target motif, X-*p*(S/T)-P (Figure 5). The vasopressin-induced decrease in ERK activity provides an explanation for the general decrease in the phosphorylation of proline-directed sites in our current dataset that includes kinases like Src (S74), Mylk (S355, S935), Mark3 (S583) and many others (Table 1). Based on the analysis shown in Figure 6, 39% (Sub-group-1, n= 121) of phosphorylation changes in response to vasopressin in this study could be due to downregulated Mapk3/ERK1 activity.

### Myosin Light Chain Kinase (Mylk)

Mylk is a calcium/calmodulin-dependent protein kinase (Figure 8) named for its ability to phosphorylate myosin regulatory light chains (Myl9, Myl12a and Myl12b), thereby activating conventional myosins Myh9 and Myh10 in a variety of cell types. Aside from its canonical target, Mylk has been shown to phosphorylate a variety of other substrates in mpkCCD cells (77). In prior studies in collecting duct cells, Mylk has been shown to be regulated by vasopressin in three ways: 1) long-term vasopressin exposure triggers a greater-than 50% decrease in the total cellular abundance of Mylk in mpkCCD cells (29); 2) vasopressin increases intracellular calcium causing a Mylk-dependent short-term increases in osmotic water permeability in isolated perfused collecting ducts presumably through regulation of AQP2 trafficking (22); 3) short-term vasopressin exposure triggers a large decrease in phosphorylation at S355 of Mylk, a putative ERK phosphorylation site, in the nucleus of mpkCCD cells (78). In the present study, we confirmed the decrease at S355 and also found a similar decrease in phosphorylation at S935, also a likely ERK target. In general, ERK mediated phosphorylation of Mylk is associated with increased activity (79, 80), so decreases in phosphorylation in the present study would quite possibly be associated with decreased catalytic activity, at least in the nucleus. Isobe et al. proposed that the effect of vasopressin on Mylk activity may be different in the nucleus versus the cytoplasm, with Ca-calmodulin-stimulated increases in cytoplasm and phosphorylation associated decreases in the nucleus (77).

### Summary

In conclusion, we have identified 25 protein kinases downstream from PKA in kidney collecting duct whose phosphorylation is altered in response to vasopressin. We have linked these kinases to the downstream substrates based on target sequence specificities and direction of change. We have also mapped them to hypothesized cellular and physiological functions (Figure 3). Little is known about the general physiological role of most of these protein kinases (e.g. Erbb2, Lmtk2, Map3k11, Nrbp2, Map3k2, Map3k5, Map3k3, Map4k5, Mink1, Taok1, Nek4, Aak1, Peak1, Srpk1, Eif2ak2 and Scyl2) in collecting duct. However, several downstream kinases (Sik2, Camkk2, Cdk18, Prkd2, Mapk3 and Mylk) seem to have important physiological roles.

## ACKNOWLEDGEMENTS

The work was primarily funded by the Division of Intramural Research, National Heart, Lung, and Blood Institute (project ZIA-HL001285 and ZIA-HL006129, M.A.K.). Protein mass spectrometry was done in the NHLBI Proteomics Core Facility (Marjan Gucek, Director). The authors thank Angel Aponte and Guanghui Wang of the NHLBI Proteomics Core Facility for mass spectrometry assistance. We thank Kavee Limbutara and Lihe Chen for helpful discussions.

## AUTHOR CONTRIBUTIONS

MAK and AD designed research; AD, CRY, VR and MAK performed bioinformatics analysis; AD, KS and MAK performed cytoscape analysis; AD did webpage generation; CLC provided useful discussion of data interpretation; AD and MAK wrote the manuscript. AD, CRY, VR, CLC and MAK edited the manuscript. All authors approved the final version of the manuscript.

## Supplemental Information

### Index

#### Supplemental Methods

All Supplementary Materials except Methods can be found at https://hpcwebapps.cit.nih.gov/ESBL/Database/PKA-Network-Supps/

**Supplemental Table 1.** Vasopressin responsive phosphopeptides in mpkCCD renal epithelial cells

**Supplemental Table 2.** Quantitative proteomics in mpkCCD cells following short-term dDAVP treatment

**Supplemental Table 3.** Kinase members from different branches of the Manning kinase dendrogram.

**Supplemental Table 4.** Rule-based classification of vasopressin-regulated phosphorylation sites

**Supplemental Table 5.** Normalized protein abundances of vasopressin-regulated substrates in differentially centrifuged fractions.

**Supplemental Spreadsheet 1.** Gene ontology (GO) and association analysis of vasopressin responsive phosphoproteins in mpkCCD renal epithelial cells

**Supplemental Spreadsheet 2.** Identification of class I PDZ ligand motif in upregulated phosphoproteins in mpkCCD renal epithelial cells

### SUPPLEMENTAL METHODS

The raw data used for this paper were reported as control experiments in our prior publication about PKA-independent signaling in collecting duct cells (1). The cell culture protocols, proteomics, phosphoproteomics sample preparation, LC-MS/MS analysis, and raw data analysis parameters remained identical as reported earlier. For the reader’s convenience, we present the detailed methodology below.

#### Reagents

Unless mentioned otherwise, all chemicals and reagents except the one used for proteomics sample preparation were purchased from Sigma. The reagents, equipment and software used for proteomics sample preparation and LC-MS/MS analysis were purchased from Thermo Fisher Scientific unless stated otherwise.

#### Cell Lines

Three clones of mpkCCD mouse kidney collecting duct cell lines, derived from *mpkCCD* clone 11-38 (mpkCCD_C11-38_) were used for this study (2). All experiments were performed with passages from 10 to 20.

#### Cell Culture: General Aspects

Cells were maintained in complete medium containing DMEM/Ham’s F-12 medium (DMEM/F-12), 2% (v/v) FBS plus supplements (5 μg/mL insulin; 50 nM dexamethasone; 1 nMtriiodotyrosine; 10 ng/mL epidermal growth factor; 60 nM sodium selenite; 5 μg/mL transferrin) at 37° and 5% (v/v) CO_2_. For all experiments, cells were seeded onto permeable membrane supports (Trans-well, 0.4 μm Polyester membrane; cat. no. 3450; Corning Costar) with complete media for 4-7 days to allow formation of a confluent epithelial monolayer with cell polarization. Then, the media were changed to serum-free simple media (DMEM/F12 containing 50 nM dexamethasone, 60 nM sodium selenite, and 5 μg/mL transferrin) for 3 days to induce cellular differentiation and to ensure complete polarization. The media used on the basolateral side was identical with the apical media except that it contains 0.1 nM of the vasopressin analog dDAVP. Transepithelial resistances were measured by voltmeter (EVOM2, WPI). The apical and basolateral media were changed daily.

#### Stable-Isotopic Labeling with Amino Acids in Cell Culture (SILAC) and dDAVP Stimulation

The cells were labeled by growing in complete medium containing either heavy (^13^C_6_^15^N_4_ arginine and ^13^C_6_ lysine) and light (^12^C_6_^14^N_4_ arginine and ^12^C_6_ lysine) amino acids. The cells were cultured for 17 days (five passages) to reach >99.9% labeling (2, 3). Heavy- or light-labeled cells were then grown on separate 6-well Transwell plates (containing labeled amino acids) for 7 days in complete and for 3 days in simple medium in the presence of dDAVP (0.1 nM) in basolateral media. On 10^th^ day, following a 2 h dDAVP wash-out period, light-labeled and heavy-labeled plates were treated with 0.1 nM dDAVP or vehicle, respectively, for 30 min. The cells were PBS-washed and stored immediately at −80°C until further processing.

#### Sample Preparation for Total Proteomics and Phosphoproteomics

The cells were thawed on ice and treated with urea buffer (8 M urea, 50 mM Tris·HCl, 75 mM NaCl, 1× Halt protease and phosphatase inhibitors), scraped and sonicated (duration: 2 min, pulse/pause: 2 sec/2 sec, output level = 1) to solubilize proteins, and then equal amounts of heavy- and light-labeled protein extracts were mixed (total, 2.5 ± 0.4 mg). The mixed samples were reduced with 20 mM DTT for 1 h at 37 °C, and then alkylated with 40 mM iodoacetamide for 1 h at 25 °C in the dark. The samples were diluted with 20 mM triethylammonium bicarbonate buffer (pH 8.5) to 1 M urea, and then digested with trypsin/LysC (Promega) (1:20 wt/wt) overnight at 37 °C. The peptides were desalted using hydrophilic–lipophilic–balanced extraction cartridges (Oasis, 1 cc, 30 mg). The peptide sample was divided into three parts: total peptide analysis (4%), heretofore referred as ‘Total’ and phosphopeptide enrichments (48% × 2) using Fe-NTA and TiO_2_ columns. The enrichment protocols were as described in the manufacturer’s instructions with minor modifications. The enriched peptides from Fe-NTA columns were desalted using graphite columns. All three samples (Total, TiO_2_, Fe-NTA) were fractionated in a 96-well plate with off-line high-pH reverse-phase chromatography (Agilent 1200 HPLC system) using a XBridge BEH C18 column (Waters; 130Å, 5 μm, 2.1 mm X 30 mm). Eluted samples were pooled to 24 (Total), 12 (TiO_2_) and 12 (Fe-NTA) fractions, vacuum-dried and stored at −80 °C. The dried peptides were reconstituted with 0.1% (v/v) formic acid (FA) in LC-MS–grade water (J.T. Baker) before mass spectrometry analysis.

#### Nano Liquid Chromatography-Tandem Mass Spectrometry (nLC-MS/MS)

Reversed-phase capillary HPLC separations were performed using a Dionex UltiMate 3000 RSLCnano system coupled in-line with an Orbitrap Fusion Lumos Tribrid mass spectrometer. Approximately 1.2 μg of peptides (calculated from digested cell lysate) were loaded onto the trapping column (PepMap100, C18, 75 μm × 2 cm), for 8 min at a flow rate of 5 μL/min with buffer A (0.1% (v/v) FA). The trapped peptides were fractionated on an analytical column (PepMap RSLC C18, 2 μm, 100Å, 75 μm i.d. × 50 cm) using a gradient of buffer B (ACN, 0.1% (v/v) FA) as follows: 4-22% for 83 min; 22-32% for 17 min; 32-90% for 3 min; followed by 90% buffer B for 7 min. The method duration was 120 min. Phosphopeptide-enriched samples were injected twice for higher-energy collisional dissociation (HCD) and electron-transfer/higher-energy collision dissociation (EThcD) fragmentation while ‘Total’ samples were analyzed by HCD fragmentation only.

#### HCD Fragmentation

MS(/MS) data were acquired on an Orbitrap Fusion as follows: All MS1 spectra were acquired over m/z 375-1500 in the orbitrap with a resolution of 120,000 (FWHM) at m/z 200; automatic gain control (AGC) was set to accumulate 4 × 10^5^ ions, with a maximum injection time of 50 ms. The intensity threshold for fragmentation was set to 25 000 and included charge states +2 to +5. A dynamic exclusion of 15 s was applied with a mass tolerance of 7 ppm. Data-dependent tandem MS analysis was performed using a top-speed approach and the dynamic parallelization using ADAPT technology was activated. The isolation window (m/z) was set at 1.6. HCD collision energy was set at 30%. MS2 spectra were acquired with Orbitrap resolution of 30 000 and a fixed first m/z of 110. AGC target was set to 5 × 10^4^, with a maximum injection time of 60 ms. The cycle time was 3 sec.

#### EThcD Fragmentation

Most of the MS(/MS) parameters for EThcD was identical as HCD fragmentation except the following: The charge states for fragmentation range from +3 to +7. ETD calibrated charge-dependent parameters and ETD supplemental activation (SA) were applied for data-dependent tandem MS. SA collision energy for EThcD was set at 15%. AGC was set to 5 × 10^4^, with a maximum injection time of 120 ms.

#### MS Raw Data Analysis

Raw data were analyzed using Proteome Discoverer 1.4 (version 1.4.0.288) in conjunction with two search engines; Mascot and SequestHT. The protein database used for SequestHT was the mouse Swiss-Prot (downloaded on April 16, 2017). The Mascot search engine and database was maintained by NIH Mascot server (https://biospec.nih.gov). The search criteria for all HCD raw files were set as follows: 1) precursor mass tolerance = 10 ppm, 2) fragment mass tolerance = 0.02 Da, 3) enzyme specificity was set as trypsin with two missed cleavages allowed. Carbamidomethylation of cysteine (C+57.021 Da) was set as a fixed modification. Variable modifications were as follows: a) isotope labeling of lysine (K+6.020 Da) and arginine (R+10.008 Da), b) oxidation of methionine (M+15.995 Da), c) deamination on glutamine and asparagine (N, Q+0.984 Da), d) phosphorylation of serine, threonine and tyrosine (S, T, Y+79.966 Da) and e) acetylation of protein N-terminal. The FDR was calculated by the target-decoy algorithm.

The search parameters for raw data files generated through EThcD fragmentation were kept identical except following modifications: 1) precursor selection: ‘use MS(n - 1) precursor’ instead of ‘use MS1 precursor’; 2) maximum precursor mass: 8000 Da instead of 5000 Da; 3) total intensity threshold: 0 instead of 1000; 3) maximum missed cleavage sites: 3. In addition, for SequestHT search, ‘c’ and ‘z’ ions were used for spectral matching instead of ‘b’ and ‘y’ ions that was used for HCD raw files.

Relative quantification of peptides and phospho-peptides was performed using the Quantification Module within Proteome Discoverer for SILAC data, which calculates relative peptide abundance ratios from light and heavy channels using the areas under curve for reconstructed MS1 ion chromatograms corresponding to each peptide. Missing values in a SILAC channel were replaced by the minimum intensity detectable for the run. The maximum ratio is set to 1000 and the minimum ratio is set to 0.001. Light: heavy ratios were normalized by dividing the peptide ratios by the median peptide ratio for each channel. The probabilities of the phosphorylation site localizations were calculated based on the given MS2 data using the *PhosphoRS 3.0* module within Proteome Discoverer. The following data reduction filters was used; peptide confidence: high, peptide rank = 1. The FDR was calculated by the target decoy PSM validator. The FDR was set <0.01.

#### Data Integration across Biological Replicates

For ‘Total’ proteomics, the SequestHT and Mascot search results of three SILAC experiments (biological replicate = 3) were integrated in Proteome Discoverer after disabling the ‘protein grouping’ option. The ratio (dDAVP/vehicle) obtained by SequestHT was used when available. Otherwise the ratio generated by Mascot was taken. The proteins that were quantified in all three biological replicates were used for further analysis. Paired *t* tests were used to calculate *p* values for comparison of log_2_[dDAVP/vehicle] ratios versus log_2_[1] (null hypothesis). The median of log-ratios was reported along with corresponding *p* value.

For phosphoproteomics, the phosphopeptides having an area (MS1 scan) of at least 1.0E7 in each of the replicate cell clones along with a corresponding median ratio calculated from TiO_2_ and Fe-NTA results were considered for further analysis. The data files obtained from HCD and EThcD fragmentation were combined. For duplicate peptides, the ones with lowest standard deviation among three replicate clones were selected. Co-efficient of variation (CV) was calculated to provide an estimate of the biological variation among three independent clones. Phospho-peptides were considered changed in abundance if they met the following criteria:ļ Log_2_(dDAVP/vehicle) |>0.4 and (-log_10_(CV))>0.4 (1). Amino acid sequences of phosphopeptides were centralized around the phosphorylation site using PTM Centralizer (https://hpcwebapps.cit.nih.gov/ESBL/PtmCentralizer/).

